# AlbaTraDIS: Comparative analysis of large datasets from parallel transposon mutagenesis experiments

**DOI:** 10.1101/593624

**Authors:** Andrew J. Page, Sarah Bastkowski, Muhammad Yasir, A. Keith Turner, Thanh Le Viet, George M. Savva, Mark A. Webber, Ian G. Charles

## Abstract

**Background:** Bacteria have evolved over billions of years to survive in a wide range of environments. Currently, there is an incomplete understanding of the genetic basis for mechanisms underpinning survival in stressful conditions, such as the presence of anti-microbials. Transposon mutagenesis has been proven to be a powerful tool to identify genes and networks which are involved in survival and fitness under a given condition by simultaneously assaying the fitness of millions of mutants, thereby relating genotype to phenotype and contributing to an understanding of bacterial cell biology. A recent refinement of this approach allows the roles of essential genes in conditional stress survival to be inferred by altering their expression. These advancements combined with the rapidly falling costs of sequencing now allows comparisons between multiple experiments to identify commonalities in stress responses to different conditions. This capacity however poses a new challenge for analysis of multiple data sets in conjunction.

**Results:** To address this analysis need, we have developed ‘AlbaTraDIS’; a software application for rapid large-scale comparative analysis of TraDIS experiments that predicts the impact of transposon insertions on nearby genes. AlbaTraDIS can identify genes which are up or down regulated, or inactivated, between multiple conditions, producing a filtered list of genes for further experimental validation as well as several accompanying data visualisations. We demonstrate the utility of our new approach by applying it to identify genes used by *Escherichia coli* to survive in a wide range of different concentrations of the biocide Triclosan. AlbaTraDIS automatically identified all well characterised Triclosan resistance genes, including the primary target, *fabI*. A number of new loci were also implicated in Triclosan resistance and the predicted phenotypes for a selection of these were validated experimentally and results showed high consistency with predictions.

**Conclusions:** AlbaTraDIS provides a simple and rapid method to analyse multiple transposon mutagenesis data sets allowing this technology to be used at large scale. To our knowledge this is the only tool currently available that can perform these tasks. AlbaTraDIS is written in Python 3 and is available under the open source licence GNU GPL 3 from https://github.com/quadram-institute-bioscience/albatradis.

## Background

Bacteria can evolve and adapt very rapidly to a wide range of challenging conditions, for example exposure to an antimicrobial. The ability of bacteria to survive antimicrobial stress is of major importance because, if current trends continue, it is predicted that by 2050 10 million people will die annually due to anti-microbial resistance (1). Despite its importance, interactions between antimicrobials and bacteria are only partially understood and most knowledge has been gained from a relatively simple set of laboratory culture conditions. Whilst the primary modes of action for most anti-microbials are known (2,3), secondary modes of action are either less well known, or not explored at all. Mechanisms of antimicrobial action and resistance in bacteria are complex and often vary depending on growth phase and/or concentration of the antimicrobial applied. A notable example of this has been described for the biocide Triclosan. Triclosan is a canonical fatty acid inhibitor although against *Escherichia coli* it exerts a bacteriostatic effect at low concentrations but is bactericidal at high concentrations (4). Additionally, understanding bacterial genotype-phenotype associations in different environments and stress conditions might help to maximise the promising health benefits from symbionts that are part of the human microbiome.

Transposon mutagenesis is an empirical tool that can provide insights into mechanisms involved in survival and fitness by simultaneously assaying the role of many genes under different conditions. This works by testing millions of mutants of a bacterial strain in parallel under various growth conditions. In this way information on gene essentiality, gene function and genetic interactions under different growth conditions can be collected (5,6). There are a number of techniques which are based on transposon mutagenesis and these include: transposon sequencing (Tn-seq) (7); high-throughput insertion tracking by deep sequencing (HITS) (8); insertion sequencing (INseq) (9); and transposon-directed insertion-site sequencing (TraDIS) (6).

Transposon mutagenesis involves randomly inserting a transposon into a bacterium to produce a mutant. On average there is a single insertion of the transposon sequence in each bacterial cell. Some of these random insertions will disrupt gene function or expression, which could potentially lead to changes in fitness (10). The mutant library can then be grown in different conditions. In some cases, the insertion will disrupt systems that are essential for life, and the bacterium will not grow (11). The corresponding gene can thus be identified as being essential for life under the given conditions by its absence from the mutant pool after growth. Likewise, when a single gene supports many insertions and growth still occurs, that gene can be considered as non-essential for growth in that condition.

Genes can be essential under one growth condition and non-essential in another. For example, bacteria may be able to expel low concentrations of antimicrobials relatively easily, but at high concentrations, above the minimum inhibitory concentration (MIC), may require different detoxification mechanisms, regulated by a different set of genes, that only become essential at high concentrations of the antimicrobial.

After exposure of the mutant library to any given condition, mutants are recovered and the transposon and a small region of genomic material from mutants are extracted and subjected to next-generation sequencing (12). The resulting sequence reads contain a short segment of the transposon and at least 45 bases of the genome adjacent to the insertion. These reads are aligned to an annotated reference genome, which allows the identification of the position at which the transposon was inserted and the insertions to be associated with specific genes and their functions. The primary output is a table of the frequencies of insertions at each base in the reference genome. Results from test conditions are compared with controls to identify conditionally important genes.

To date, one major barrier to the adoption of transposon mutagenesis for mechanistic studies has been the complex nature of the protocols and the need for non-standard sequencing instrument setups (12). These issues have been incrementally overcome which, in conjunction with the rapidly falling costs of genome sequencing, has made transposon mutagenesis an increasingly cost-effective method for screening millions of mutants simultaneously under a large number of different conditions (5,13–16).

A limitation of the traditional TraDIS approach is, that essential genes cannot be effectively assayed, as mutants with insertions in them will not grow. A recent modification of the TraDIS protocol (17) (TraDIS+) allows the conditional fitness of all genes in the genome to be assayed simultaneously, including essential genes. This methodology uses a transposon with an outward directed inducible promoter allowing the impact of transcription alteration of each gene to be assayed as well as gene inactivation. By comparing induced and uninduced conditions a better ‘signal-to-noise’ ratio is achieved to identify genes where expression changes contribute to conditional survival. Additionally, it is a suitable approach to identify where ‘knock-down’ of expression of a gene can influence survival. Incorporating the ability to alter expression of all the genes of an organism in one experimental condition in a controlled manner promises to be hugely powerful, as applying changes to all genes in a genome without prior knowledge about function has the potential to uncover a large number of new genotype-phenotype relationships.

Analysis of the large-scale highly complex data resulting from experiments using transposon mutagenesis can be a considerable challenge; analysis involves tens of millions of data points (each corresponding to a physical bacterium), with controls and multiple replicates. The interpretation of these data is thus complicated. Previous work has focused on manually interpreting insert site patterns by comparing mutants with controls (18) or by looking for simple signals that indicate whether a gene is essential for the survival of a bacterium (16), or for its evolutionary fitness using tools such as Bio-TraDIS (12). However, modes of action and any commonalities between different growth conditions are not computationally identified within the existing Bio-TraDIS toolkit, and results must be manually analysed. This is time consuming and limits the number of conditions that can be compared. While the Bio-TraDIS toolkit identifies essential and non-essential genes as well as performs comparison between one condition and control, it has little functionality for filtering, prioritising and cross conditional comparison. In order to evaluate the putative genes identified by the Bio-TraDIS toolkit, a visualisation tool, such as Artemis (19), must be used to compare multiple replicates for a condition against controls. This requires prior knowledge and experience to judge which inserts are most likely to be biologically significant. Therefore, visualising all of the information from more than a single condition becomes impractical due to the volume of information.

To address these issues, we present AlbaTraDIS, a software for rapid large-scale comparative analysis of TraDIS experiments that predicts the impact of inserts on nearby genes. It uses the statistical methods published in the Bio-TraDIS toolkit as a foundation. To our knowledge this is the only tool currently available that can perform these tasks. AlbaTraDIS is written in Python 3 and is available under the open source licence GNU GPL 3 from https://github.com/quadram-institute-bioscience/albatradis.

## Implementation of AlbaTraDIS

In the main AlbaTraDIS workflow (albatradis script), as illustrated in Figure 1, we extend the Bio-Tradis functionality to identify and analyse signals from data generated by the TraDIS+ method, which includes determining putative alteration of transcription of genes in the forward or reverse complementary directions. The input to the albatradis script are insert site plots along with the annotated reference genome in EMBL format (20). The insert site files contain the number of insertions on the forward and reverse strands, at each base in the genome. One or more growth conditions, and a matching number of controls, are required as input, with a minimum of two replicates recommended to account for experimental variation. To generate the insert site files, sequence reads generated using Illumina sequencing, are aligned to a reference genome using Bio-Tradis.

**Figure 1:**
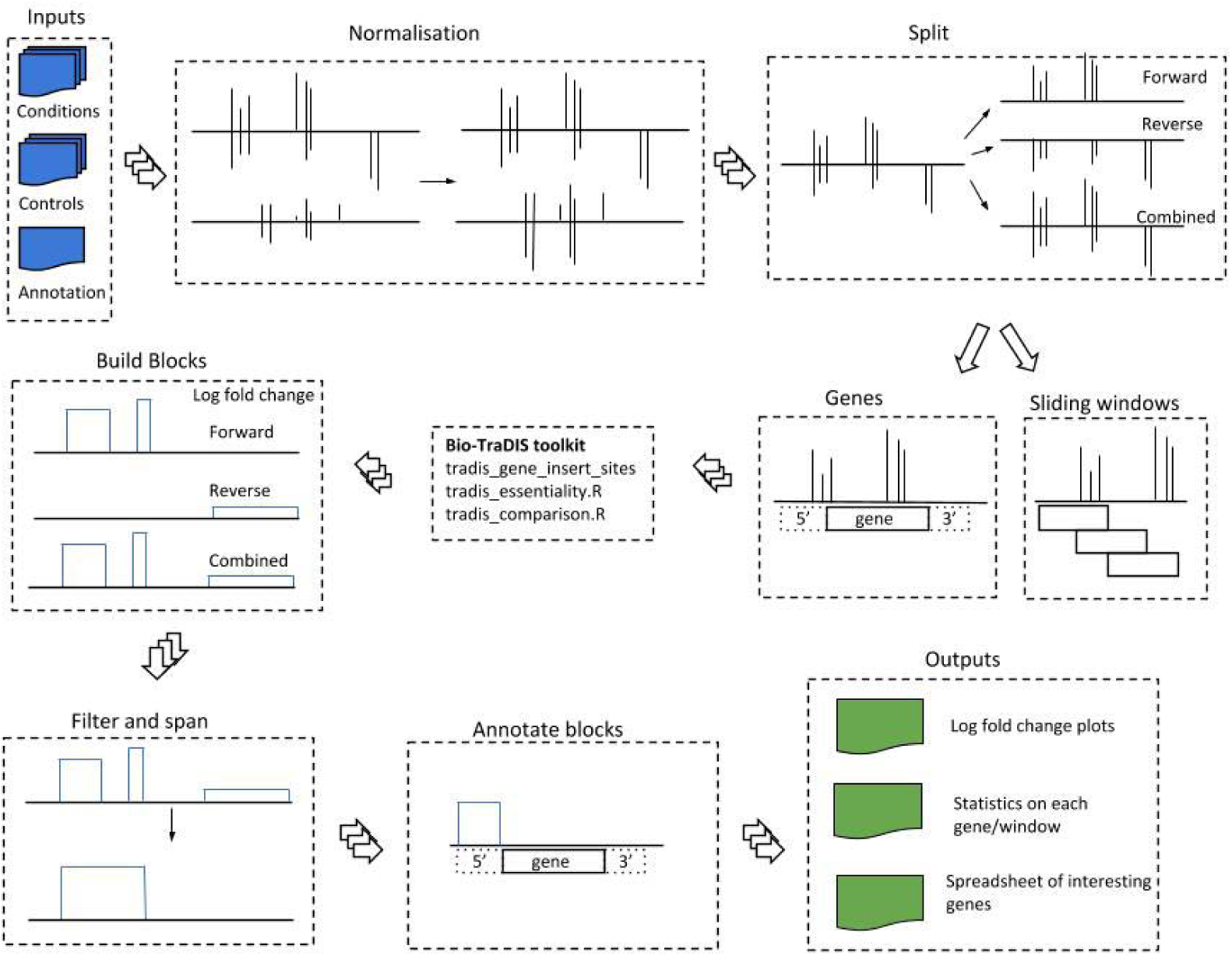
The underlying method for AlbaTraDIS. The inputs are insert site plots, with a frequency count of the insertions at each base in the genome for a condition and controls and the annotated genome in EMBL format. The abundance of inserts are normalised and the plots split into forward strand, reverse strand and combined strand insertions. Essentiality and differential abundance is assessed using sliding windows or a per gene option. The height of the log fold change plot indicates the log fold change difference in insertions between the conditions and controls. The list of significant genes is compiled using user definable values of corrected p value (q-value), logCPM and logFC.

The first step in the albatradis workflow is to apply normalisation in order to provide a more consistent analysis in the presence of natural experimental variation, but this option can be disabled if it is not desired. Each input file is normalised by the ratio of the number of insertions in the input file to the maximum number of insertions across all files.

In order to screen the genome for different signals, by default, a reference-free sliding window is used. The window size defaults to 50 bases, as this was found experimentally to be the minimum window size where a signal could be detected with an insertion site density of one insertion every ten bases. This can be increased, but the boundaries of an identified mechanism become poorly defined, or may be missed entirely, if multiple mechanisms are present within one window, cancelling each other out. Alternatively, there is an option for an annotated reference-guided analysis. Each of the annotated genes and features are then treated as windows.

Genes and Windows are annotated with their essentiality. An essential gene is a gene which has no or very few insertions (no data points) as without the functioning gene, the bacteria do not survive, and thus are not present in the resulting sequencing data from that particular experiment (See Figure 2B). Essentiality analysis is performed using the method as implemented in Bio-Tradis (tradis_essentiality.R). A threshold value for the number of insertions within essential genes is estimated using the observed bimodal distribution of insertion sites over genes when normalized for gene length (5).

**Figure 2:**
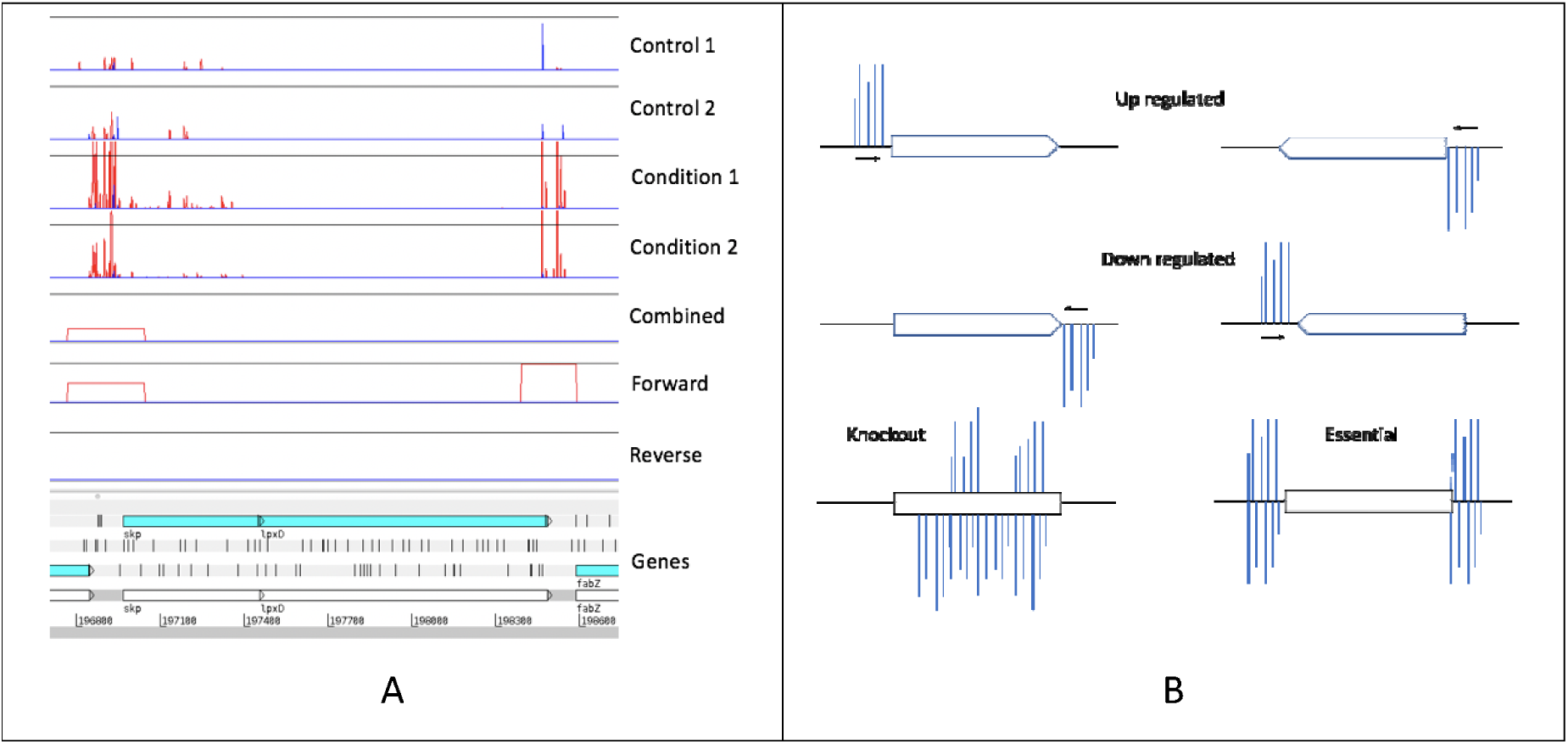
A) The top four lines are the insertion sites in controls and under treatment conditions, where red lines are insertions in the forward direction and blue lines are insertions in the reverse direction, with the height corresponding to the number mapped reads identified for this site. The next three lines correspond to the signal identified by AlbaTraDIS using a sliding window of 50 bases and an interval of 25 bases, with the height corresponding to the log fold change between the treatments and controls. The bottom section shows the genes as found in the reference genome, with the forward reading frames of translation. B) The pattern of insertions around a gene that Imply transcriptional augmentation, in the forward or reverse complementary direction. The shape of the gene indicates the direction, with the 5’ at the beginning (flat end) and the 3’ prime at the pointed end. Insertions on the forward strand are above the line and insertions on the reverse strand are below the line.

The log FC of each window, or gene, is overlaid onto the bases of the genome, producing plot files for analysis of the forward, reverse and combined data, and visualisation in applications such as Artemis (19).

If the sliding window option is used, short gaps are spanned automatically. This shows where there is a strong increase or decrease of insertions in any part of the genome, and whether it is in a single direction, or in both directions. This translates multiple signal spikes into clearly delineated blocks with putative modes of action (See Figure 2A). Any regions of the genome with blocks or genes above pre-defined levels (as previously noted) are selected as loci that may have a putative role in sensitivity to the test conditions. Putative changes in the numbers of mutants with insertions upstream or downstream of genes which may alter transcription are strong indications that those genes are important in bacterial survival under test conditions and also allows inferences about the importance of essential genes.

In order to identify insertions that may alter gene transcription as well as knockouts, the insertions are divided into the forward and reverse inserts, giving three streams for analysis (forward, reverse and combined). The aim is to identify significant changes in each sliding window or gene between condition and control as described in (5) (See Figure 2B). This analysis is based on methodology used for differential expression analysis as implemented in edgeR (21), as the data is given as insertion counts per gene or genetic region and can therefore be modelled by a negative binomial distribution. Therefore, the next step in the albatradis workflow is calling the Bio-TraDIS toolkit (tradis_comparison.R) to perform comparison of insertion abundances between control and condition. This comparison comprises a normalisation of trimmed mean of M values (TMM) (22) and the calculation of distribution parameters based on tag-wise dispersion estimates. The resulting distributions for condition and control are then compared using an adopted exact test. P values are corrected for multiple testing using the Benjamini-Hochberg method (23). A list of all significant genes is produced. The user can specify parameters that mark significance, but as a default a corrected P value (Q value) of < 0.05, an absolute log fold change (log FC) of > 1, and an absolute log count per million (log CPM) > 8 are considered significant. The produced list also contains a summary of each statistically-significant gene, its classification (up/down regulation, knockout), its coordinates, its maximum log FC, whether there is increased or decreased expression, the direction of the signal (forward/reverse strain, or both) and the upstream gene.

### Multiple condition comparison

The albatradis main workflow compares replicates of one control and one condition. Often there are many different conditions and/or timepoints. Aiming to give a more complete picture of what happens, it is of interest to compare the different conditions/timepoints in order to identify commonalities. The albatradis-presence_absence script summarises and performs comparative analysis of the outputs from the albatradis workflow. The impact of each test condition on each gene can be observed. Changes in essentiality of genes are compared with the control (i.e., where essential genes become non-essential and where non-essential genes become essential). All of these methods are designed to allow scaling up and automation of the TraDIS analysis. The input to the script are multiple *gene reports*, representing various test conditions and the annotated genome (embl format). A variety of outputs is produced: the union and intersection of the genes for the test conditions which allows for further analysis of commonalities, a global heatmap of the log FC observed between the conditions and the controls and a spreadsheet representing the heatmap data. Common patterns can be represented by a tree structure, grouping common biological modes of action together. Two trees are created, one using hierarchical clustering (dendrogram) and one using the neighbour-joining method. Both trees are supplied in Newick format (http://evolution.genetics.washington.edu/phylip/newicktree.html) and can be viewed using a visualisation program like FigTree (24) (See Figure 3A).

**Figure 3:**
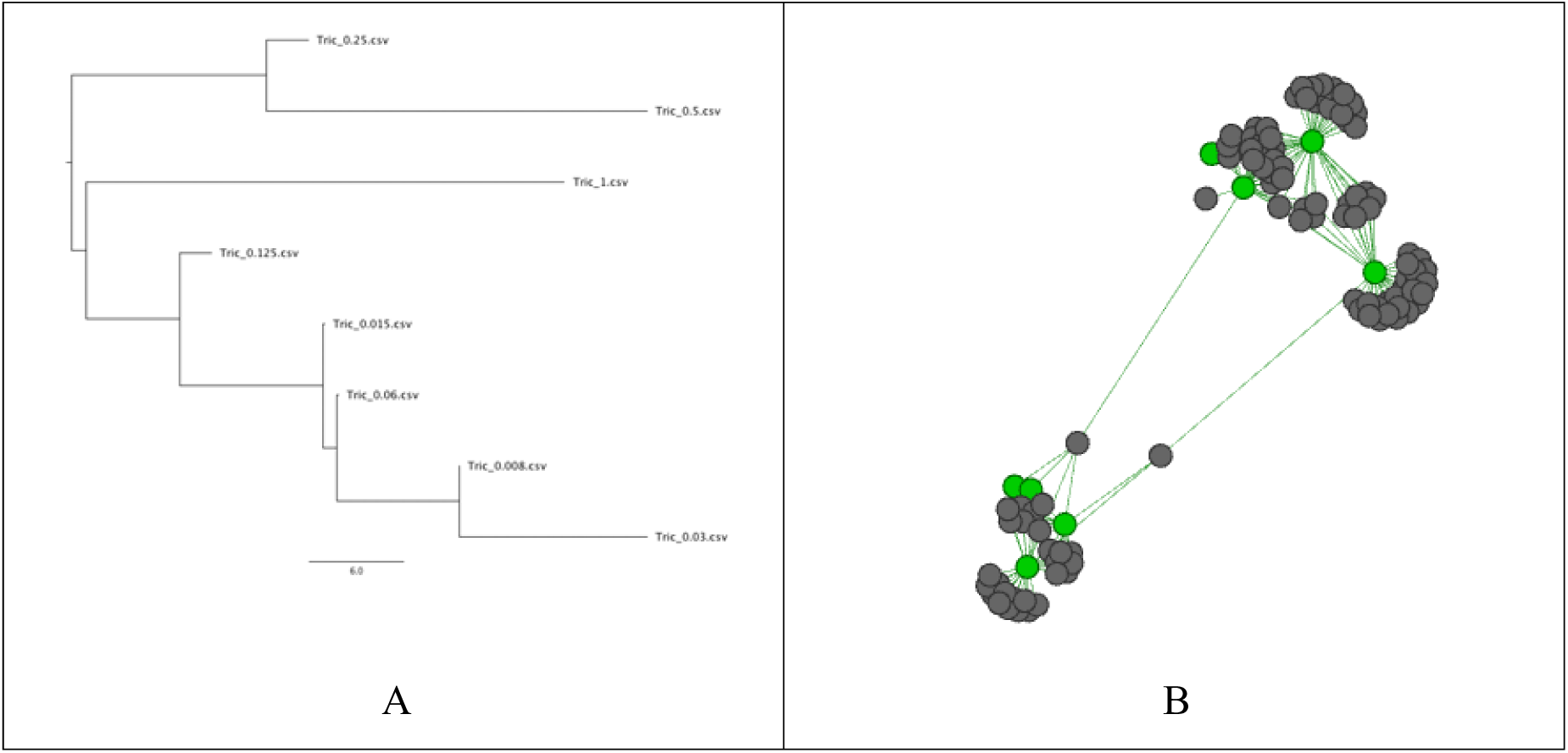
A) Neighbour joining tree of the presence and absence of genes that have significant differences in the number of insertions compared with the control after exposure to different concentrations of Triclosan. This shows how similar different conditions relate to each other based on their modes of action. B) Example network of the relatedness of different modes of action where the green nodes are different conditions (such as drug concentrations), and the grey nodes are a single gene.

A graphical representation of the collection of genes under different conditions is provided. Genes and conditions are represented as nodes in the graph. Where AlbaTraDIS has identified a link between a test condition and a gene, an edge is added, which is weighted by the number of identified connections. Figure 3B gives an example of such a network. The grey nodes represent genes and the green nodes represent test conditions. This allows for interrogation of commonalities between conditions using standard graph theory algorithms. If there are no genes in common amongst the conditions, the graph consists of several disconnected subgraphs.

## Results

### Experimental data used to evaluate usefulness of AlbaTraDIS

To evaluate the performance of AlbaTraDIS, it was used to analyse a dataset from TraDIS+ experiments of *E. coli* grown in different concentrations of the antibacterial agent, Triclosan. This showed that large scale analysis was possible and confirmed the identity of known modes of action. A full description of this dataset is given in the companion article (17); this is the first dataset of this scale to be published. We briefly summarise the experiments and the data collected.

Triclosan is an antibacterial agent that has been widely used in clinical practice and in cleaning and domestic hygiene products (25). It is known to exhibit concentration dependent effects; at low concentrations it is bacteriostatic (inhibits growth) and bactericidal (kills) at high concentrations (25,26). However, the mechanisms for these modes of action are not well understood, with only one primary target well validated (25). TraDIS was used to gain a better understanding of the consequences of exposure to Triclosan at different concentrations with E. coli BW25113 (27). This bacterium was chosen because it is well characterised laboratory strain with a fully sequenced genome. *E. coli* BW25113 is also the parent strain of the Keio collection (28), for which every gene in the genome has been systematically knocked out, allowing for subsequent experimental validation of phenotype. A library of around half a million mutants was generated from *E.coli* BW25113 using a transposon that contained an inducible outward directed promoter. The promotor allowed for enhanced expression with Isopropyl β-D-1-thiogalactopyranoside (IPTG). The mutant library was then grown for 24 hours in eight concentrations of Triclosan (from 0.008 to 1 mg/L) and in combination of three concentrations of the inducer to give a spectrum of promoter expression. There were two controls and two technical replicates, resulting in 60 individual TraDIS experiments. Table 1 provides the accession numbers for data collected and the conditions evaluated (Triclosan concentrations) for each experiment. The genome of *E. coli* BW25113 (accession number GCA_000750555.1) (27) consists of 4,631,469 bases in a single chromosome with 4,774 annotated genes.

**Table 1:** Conditions evaluated (Triclosan concentrations) and accession numbers for each experiment. The overall project accession number is PRJEB29311.

| Triclosan (mg/L) | Accession number for experiment |  |
| --- | --- | --- |
|  | Replicate 1 | Replicate 2 |
| 0.008 | ERR2854367 | ERR2854368 |
| 0.015 | ERR2854369 | ERR2854370 |
| 0.03 | ERR2854371 | ERR2854372 |
| 0.06 | ERR2854373 | ERR2854374 |
| 0.125 | ERR2854375 | ERR2854376 |
| 0.25 | ERR2854377 | ERR2854378 |
| 0.5 | ERR2854379 | ERR2854380 |
| 1.0 | ERR2854381 | ERR2854382 |
| Control 1 | ERR2854363 | ERR2854364 |
| Control 2 | ERR2854365 | ERR2854366 |

### Ability of AlbaTraDIS to identify primary modes of action

To confirm that the results from AlbaTraDIS are accurate, we used it to evaluate the Triclosan dataset for *E. coli* BW25113 as listed in Table 1. We looked for the presence of genes that are known from experimental validation to be important in the action of Triclosan, and also important in bacterial resistance to Triclosan. The primary target of Triclosan is the enzyme FabI. Mutation or over-expression of *fabI* are known mechanisms of resistance to Triclosan (25). Whilst *fabI* is essential, and therefore not assayed by traditional transposon mutagenesis approaches, inserts upstream of *fabI* at the 5’ end were clearly identified by AlbaTraDIS. An induction of *fabI* was classified as beneficial for survival when grown in Triclosan(Figure 4). Other genes known to be involved in resistance were also identified including the efflux and regulators *acrR, acrB, marR, soxS*, and many genes involved in generation of lipopolysaccharide. A number of loci not known previously to be involved in Triclosan resistance were also identified. The predicted phenotypes for a selection of these were validated by using the corresponding knockout mutants from the Keio library, growing them in different Triclosan concentrations for 24h and assessing their growth rate in comparison to the parent strain BW25113. The results showed high consistency with predictions. As previously mentioned, more details on these results and other biological outcomes as well as methodology can be found in the companion paper (17).

**Figure 4:**
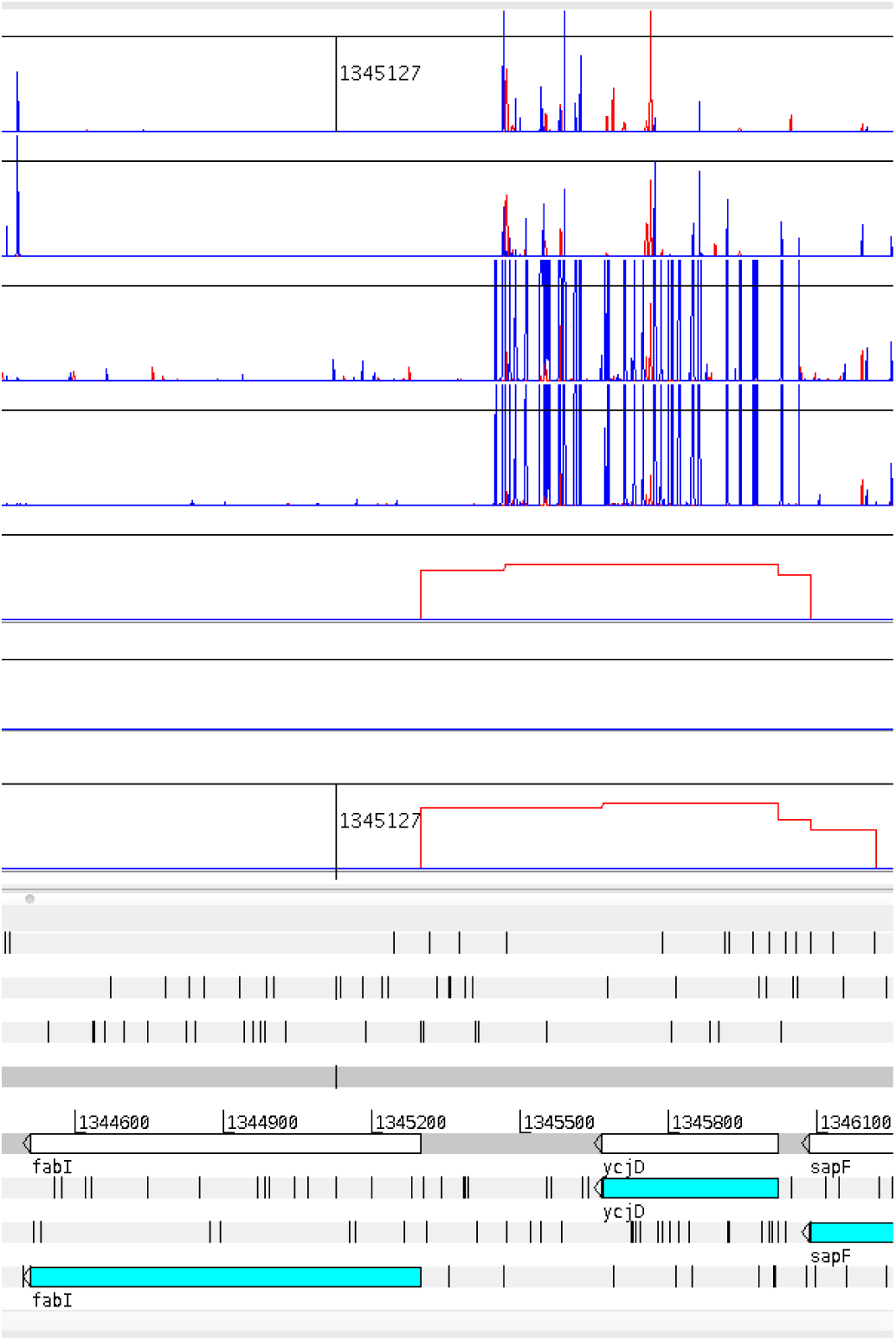
The top 4 panels show the transposon insertion sites, 2 controls and 2 for libraries grown in 0.5 mg/L triclosan. The next three lines correspond to the signal identified by AlbaTraDIS using a with the height corresponding to the log fold change between the treatments and controls. There is an increase in insertions in the promotor area (upstream) in the direction towards the gene, which indicates that up-regulation of fabI in E. coli grown in 0.5 mg/L of Triclosan might be beneficial to survival. This shows that AlbaTraDIS can identify the primary target of Triclosan.

### Computational Environment

All of the computational experiments were performed on the MRC CLIMB framework (30), using the Genomics Virtual Laboratory (v4.2) (31). The operating system was Ubuntu 16.10 LTS and the resources available were four processors and 32 Giga Bytes (GB) of memory. Resources of this scale are not required to run AlbaTraDIS, these were merely the default minimum available. AlbaTraDIS version 0.0.5 running on Python version 3.6.7 and Bio-TraDIS version 1.4.1 (12) running on Perl version 5.22.1 were used. Experimental running times and peak memory usage were measured using the time command.

### Performance of AlbaTraDIS

The scalability of AlbaTraDIS was evaluated by varying the number of test conditions included in the analysis using the data and computing resources described. As the number of test conditions increased the total running time of the main AlbaTraDIS workflow increased linearly (See Figure 5) which matched the theoretical runtime (O(n)). However, the most resource-intensive part of the process can be parallelised, and runtime is constant when the number of processors equal the number of test conditions. The condition comparisons running time, while also linear, was negligible. As an indication of the overall running time, the full dataset described previously took 74 minutes when run with a single processor. This is likely to vary with the available computing resources and datasets.

**Figure 5:**
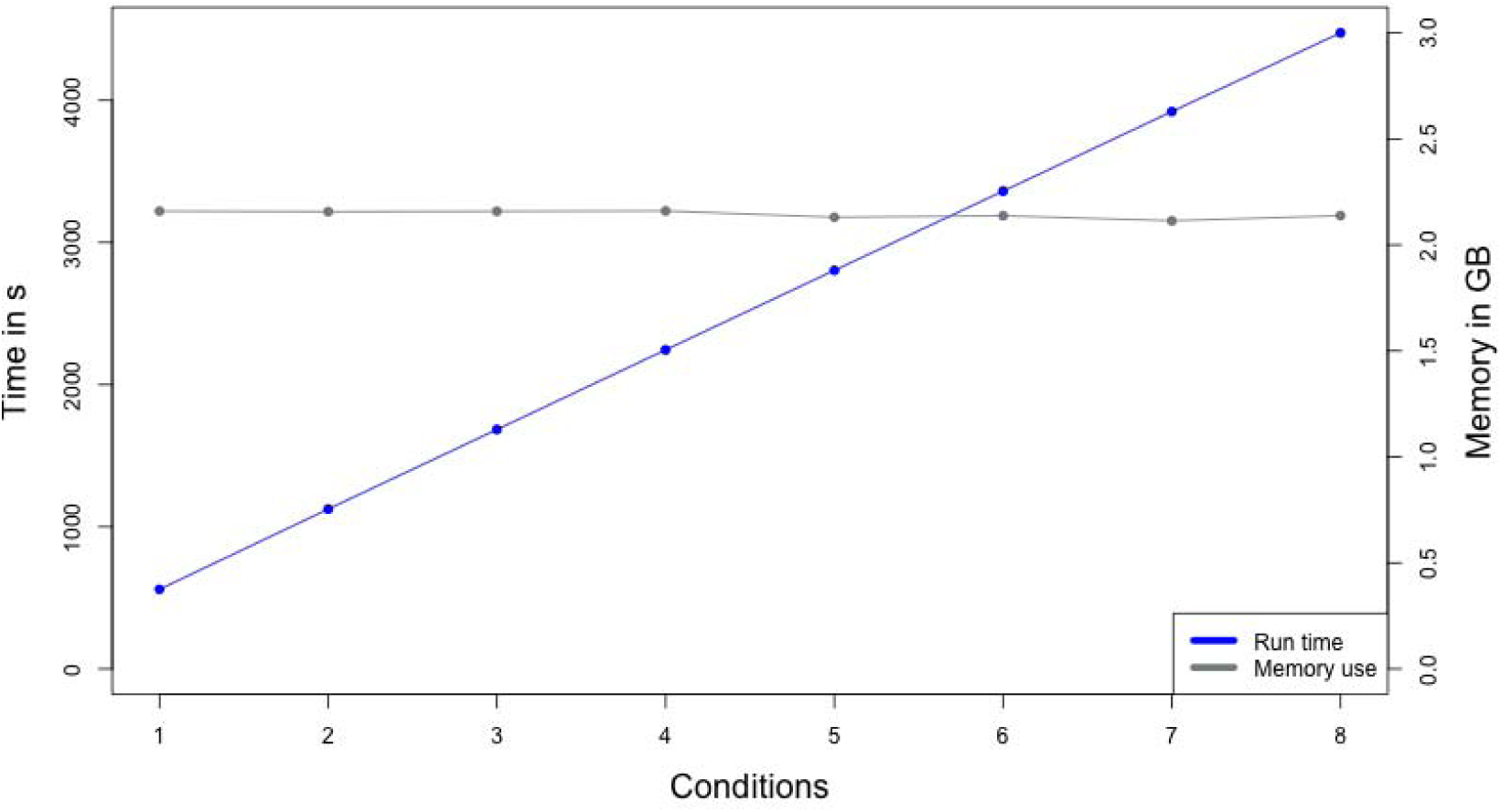
The total running time, including comparative analysis, for varying numbers of test conditions when using a single CPU (Blue) and the total memory usage in GB, including comparative analysis, for varying numbers of conditions when using a single CPU (Grey).

The total memory usage (See Figure 5) remained constant but will vary with different datasets. When 1 processor was available for each condition (n=8), the total running time was just 9.4 minutes. The memory requirement was 2.1 GB for eight conditions, which is low enough that it can be run on a standard desktop machine. We were able to achieve these results by using Cython (32) is used internally for computationally-intensive parts of the method, allowing for native C-compiled code to be used within Python.

## Conclusions

AlbaTraDIS allows the analysis of large-scale transposon insertion sequencing experiments to be performed and results compared across conditions than had previously been possible. In addition, the context of inserts in relation to local genes and their impacts can be predicted which greatly reduces the complexity of the analysis required for large data sets. Comparative analysis of the results from a range of experimental conditions allows identification of common modes of action. Known mechanisms of resistance were efficiently identified, including those where expression changes were important. AlbaTraDIS is fast, scalable and can be run on standard desktop machines.

## Availability and requirements

**Project name:** AlbaTraDIS

**Project home page:** https://github.com/quadram-institute-bioscience/albatradis

**Operating system(s):** Linux, OSX

**Programming language:** Python version 3.3+

**Other requirements:** Bio-TraDIS toolkit

**License:** GNU GPL version 3

**Any restrictions to use by non-academics:** GNU GPL version 3

The software can be installed using *conda* (33), *pip* (https://pypi.org) or as a Docker container (34).

## List of Abbreviations

GMI: Global Microbial Identifier
IPTG: Isopropyl β-D-1-thiogalactopyranoside
NGS: Next Generation Sequencing
TraDIS: Transposon Directed Insertion-site Sequencing
MIC: Minimum Inhibitory Concentration
TMM: Trimmed Mean of M values
CPM: Count Per Million
FC: Fold Change
GB: Giga Bytes

## Declarations

### Ethics approval and consent to participate

Not applicable.

### Consent for publication

Not applicable.

### Availability of data and material

The datasets generated and/or analysed during the current study are available without restriction from the European Nucleotide Archive at EMBL-EBI and accession numbers for the raw data are listed in Supplementary.

## Authors’ contributions

AJP wrote the software and wrote the manuscript.

SB contributed to the software and the manuscript.

TLV packaged the software for general use.

KAT, MY performed the microbiology experiments and interpreted the results.

GMS, MAW, IGC provided overall study design and guidance.

All authors have read and contributed to the manuscript.

## Acknowledgements

We thank Andrea Telatin and Marianne Defernez for their involvement in this project and Lars Barquist for providing the Bio-TraDIS toolkit.

